# Species- and C-terminal linker-dependent variations in the dynamic behavior of FtsZ on membranes *in vitro*

**DOI:** 10.1101/278564

**Authors:** Kousik Sundararajan, Anthony Vecchiarelli, Kiyoshi Mizuuchi, Erin D. Goley

## Abstract

Bacterial cell division requires the assembly of FtsZ protofilaments into a dynamic structure called the ‘Z-ring’. The Z-ring recruits the division machinery and directs local cell wall remodeling for constriction. The organization and dynamics of protofilaments within the Z-ring coordinate local cell wall synthesis during cell constriction, but their regulation is largely unknown. The disordered C-terminal linker (CTL) region of *Caulobacter crescentus* FtsZ (*Cc*FtsZ) regulates polymer structure and turnover in solution *in vitro*, and regulates Z-ring structure and activity of cell wall enzymes *in vivo*. To investigate the contributions of the CTL to the polymerization properties of FtsZ on its physiological platform, the cell membrane, we reconstituted *Cc*FtsZ polymerization on supported lipid bilayers (SLB) and visualized polymer dynamics and structure using total internal reflection fluorescence microscopy. Unlike *E. coli* FtsZ protofilaments that organized into large, bundled patterns, *Cc*FtsZ protofilaments assembled into small, dynamic clusters on SLBs. Moreover, *Cc*FtsZ lacking its CTL formed large networks of straight filament bundles that underwent slower turnover than the dynamic clusters of wildtype FtsZ. Our *in vitro* characterization provides novel insights into species- and CTL-dependent differences between FtsZ assembly properties that are relevant to Z-ring assembly and function on membranes *in vivo*.

## Introduction

In bacteria, the process of cytokinesis requires remodeling of the cell wall at the division site following the assembly of the multi-protein division complex termed the divisome. The tubulin homolog FtsZ polymerizes and forms a ring-like scaffold called the “Z-ring” at the incipient division site for the recruitment of the divisome. FtsZ protofilaments assemble into dynamic clusters that together form a discontinuous Z-ring (Li *et al.*, 2007; Fu *et al.*, 2010; Holden *et al.*, 2014; Bisson-Filho *et al.*, 2017; Yang *et al.*, 2017). Following assembly of the Z-ring, more than two dozen factors are recruited to the division site through direct or indirect interactions with FtsZ (Erickson *et al.*, 2010; Meier and Goley, 2014). Through the recruitment of cell wall enzymes, the Z-ring promotes local cell wall synthesis (Aaron *et al.*, 2007). In addition to their recruitment, the Z-ring also regulates the activity of these enzymes at the site of division (Sundararajan *et al.*, 2015). Recent studies of FtsZ have suggested that the dynamics of clusters of protofilaments in the Z-ring result in an apparent directional movement of clusters through treadmilling (Bisson-Filho *et al.*, 2017; Yang *et al.*, 2017)1. Moreover, the direction and speed of these clusters are correlated with the direction and speed of movement of cell wall enzymes required for cell division. Thus, it appears that the polymerization properties of FtsZ are essential for its function in local cell wall remodeling during cytokinesis (Sundararajan *et al.*, 2015; Bisson-Filho *et al.*, 2017; Yang *et al.*, 2017). However, the regulation of the assembly of FtsZ into dynamic clusters, the higher-order arrangement of protofilaments within the clusters, and the source of directional dynamic assembly of these clusters are largely unknown.

In cells, FtsZ protofilaments observed by electron cryotomography appear as slightly curved protofilaments running circumferentially along the short axis of the cell near the inner membrane (Li *et al.*, 2007; Szwedziak *et al.*, 2014). *In vitro*, FtsZ polymerizes on binding GTP into single protofilaments, straight multifilament bundles, helical bundles or toroids depending on the presence of binding factors, crowding agents or divalent cations, as observed by electro microscopy (Mukherjee and Lutkenhaus, 1999; Gueiros-Filho and Losick, 2002; Popp *et al.*, 2009; Goley *et al.*, 2010). It is unclear which of the structures of FtsZ polymers observed *in vitro* are physiologically relevant in cells, especially in the context of attachment to the membrane. Efforts to observe dynamic assembly of FtsZ polymers on a membrane have been limited to *E. coli* FtsZ (Arumugam *et al.*, 2012; Loose and Mitchison, 2014; Arumugam *et al.*, 2014; Ramirez *et al.*, 2016). In particular, purified *E. coli* FtsZ assembles into large, dynamic bundles of treadmilling protofilaments when anchored to a membrane either using an artificial membrane targeting sequence (MTS) or membrane-anchoring proteins such as FtsA and observed by total internal reflection fluorescence microscopy (TIRFM) *in vitro* (Loose and Mitchison, 2014;Ramirez *et al.*, 2016). The conservation and physiological relevance of these emergent structures of FtsZ protofilaments on membranes have yet to be demonstrated.

FtsZ polymerizes through its conserved tubulin-like GTPase domain. The GTPase domain is followed by a C-terminal tail made of an intrinsically disordered region (C-terminal linker or CTL) and a conserved peptide region (C-terminal conserved peptide or CTC) for binding membrane-anchoring proteins of FtsZ (Vaughan *et al.*, 2004; Erickson *et al.*, 2010). In *Caulobacter crescentus, E. coli* and *B. subtilis*, FtsZ requires the CTL to assemble into a functional Z-ring capable of cytokinesis (Buske and Levin, 2013; Gardner *et al.*, 2013; Sundararajan *et al.*, 2015). In *C. crescentus* cells, expression of FtsZ lacking its CTL (ΔCTL) has dominant lethal effects on cell wall metabolism leading to cell filamentation, local envelope bulging and rapid lysis (Sundararajan *et al.*, 2015). Although ΔCTL is functional for recruiting all of the known FtsZ binding proteins and directing local cell wall synthesis, it causes defects in the chemistry of the cell wall that lead to cell lysis (Sundararajan *et al.*, 2015). The CTL thus contributes to the ability of FtsZ to regulate cell wall metabolism, independent of FtsZ’s function as a scaffold for localizing cell wall enzymes. The structures formed by ΔCTL in cells appear deformed when compared to wildtype FtsZ – they are larger, brighter, and less ring-like by epifluorescence microscopy. *In vitro*, Δ CTL polymerizes into long, straight multifilament bundles with low GTP hydrolysis rates compared to single slightly curved protofilaments of wildtype FtsZ (Sundararajan and Goley, 2017). How the most variable region of FtsZ across organisms – the CTL – contributes to the higher order assembly of the Z-ring and its function in cell wall metabolism is not fully understood.

Here, we have developed an *in vitro* TIRFM-based assay to image FtsZ polymers anchored to supported lipid bilayers (SLB) through an artificial membrane tethering sequence peptide (MTS) from *E. coli* MinD (Osawa *et al.*, 2008). Unlike prior reconstitution studies of FtsZ polymerization on membrane constrained within wells placed on coverslips (Loose and Mitchison, 2014; Arumugam *et al.*, 2014; Ramirez *et al.*, 2016), we adapted a system to allow for rapid depletion and repletion experiments in a controlled environment using flow cells (Vecchiarelli *et al.*, 2016). Using our flow cell setup, we observe that whereas *E. coli* FtsZ assembles into large, dynamic bundles, *C. crescentus* FtsZ assembles into smaller dynamic clusters under identical *in vitro* conditions. Investigating the effects of the CTL on FtsZ polymerization, we observe that ΔCTL forms large networks of straight filaments on SLBs that turn over more slowly compared to the dynamic clusters formed by WT FtsZ. We conclude that the CTL is required for disrupting lateral interaction between protofilaments and promoting polymer turnover on membranes. Our study provides the first *in vitro* characterization of assembly and dynamics on SLBs of FtsZ from an organism other than *E. coli* and describes CTL-dependent regulation of FtsZ polymerization that we propose is relevant to FtsZ-mediated regulation of cell wall metabolism in cells.

## Results

### *C. crescentus* FtsZ assembles into dynamic clusters on supported lipid bilayers

To image FtsZ polymer assembly on SLBs, we adapted the flow cell setup developed for observing MinD-MinE protein oscillations on membranes (Vecchiarelli *et al.*, 2016). Specifically, we coated flow cells with SLBs composed of combinations of synthetic anionic (DOPG) and zwitterionic (DOPC) lipids (Figure 1A). To visualize FtsZ filaments anchored to the membrane, we used fluorescently labeled FtsZ fused to the membrane targeting sequence (MTS) from *E. coli* MinD, or a mixture of unlabeled (non-fluorescent) FtsZ fused to MTS and fluorescently labeled FtsZ that had no MTS to generate copolymers. FtsZ variants were incubated with GTP and flowed into the SLB-coated flow cell (~ 3 μL total volume). FtsZ polymers on the membrane were then imaged using prism-type total internal reflection fluorescence microscopy (TIRFM) (Figure1A).

**Figure 1.**
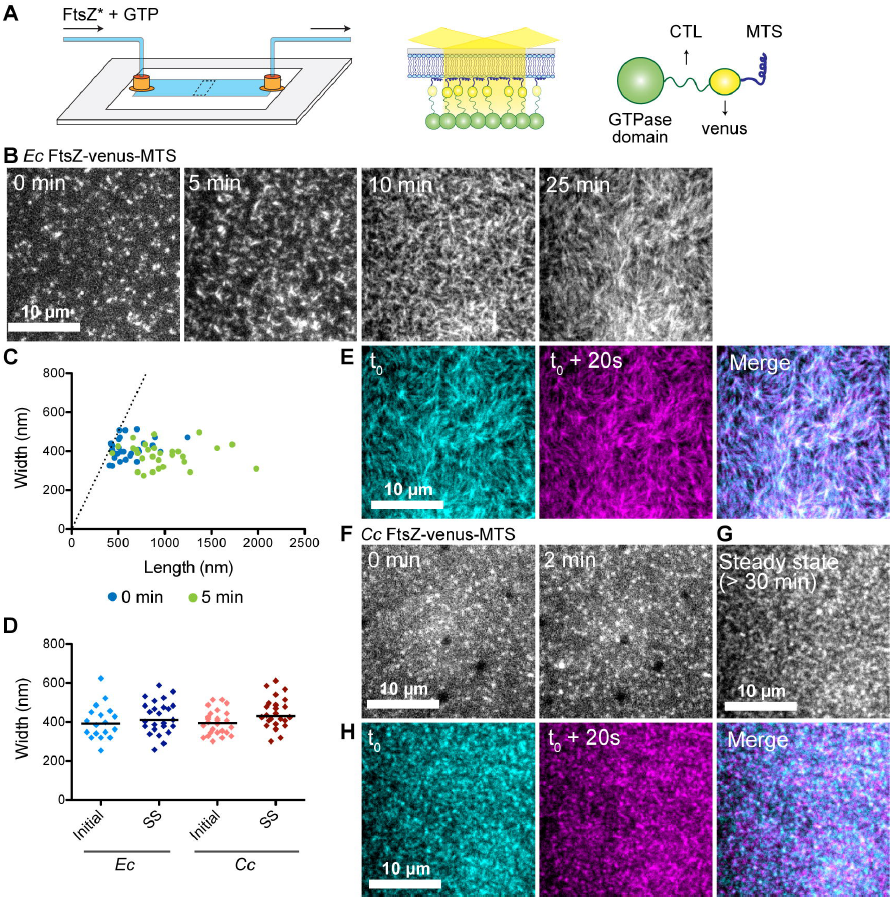
FtsZ protofilaments assemble as dynamic clusters on SLBs that form species-specific superstructures. **A.** Schematic describing the flow cell setup used for imaging FtsZ polymer assembly. FtsZ* (FtsZ-venus-MTS) incubated with GTP is flowed into the flow cell. FtsZ-venus-MTS protofilaments are recruited to the membrane through the MTS and are brought into the evanescent field of TIRF. **B.** Contrast enhanced TIRFM images showing structures formed by 2 μM *Ec* His_6_-FtsZ-venus-MTS preincubated with 2 mM GTP for 30 minutes and introduced into flow cell (at 5 μL/minute) with the SLB composed of 33% DOPG and 67% DOPC lipids. Time on the images indicates approximate time passed after the initiation of flow. **C.** Plot showing width (distance along short axis) and length (distance along long axis) of clusters formed at 0 minutes (blue) and 5 minutes (green) for experiment shown in B. Dotted line indicates the identity line (width = length). **D.** Widths of clusters or bundles formed by *E. coli* His_6_-FtsZ-venus-MTS or *C. crescentus* FtsZ-venus-MTS at initial time point (time = 0 minutes) and at steady state (time ≥ 30 minutes). Line indicates median. **E.** Individual frames and merged images showing overlay of structures formed by *Ec* His_6_-FtsZ-venus-MTS at steady state spaced 20 seconds apart (cyan – time ‘t_0_’, magenta – time ‘t_0_ + 20 seconds’, white regions in the merged image represent colocalization of signal) **F.& G.** Contrast enhanced TIRFM images showing structures formed by 18 μM *Cc* FtsZ-venus-MTS preincubated with 2 mM GTP for 30 minutes and flowed into flow cell (at 5 μL/minute) with the SLB composed of 33% DOPG and 67% DOPC lipids. Time on the images indicates approximate time passed after the initiation of flow. **G.** Steady state structures formed by *Cc* FtsZ-venus-MTS after flow was stopped. **H.** Individual frames and merged images showing overlay of structures formed by *Cc* FtsZ-venus-MTS at steady state spaced 20 seconds apart (cyan – time ‘t_0_’, magenta – time ‘t_0_ + 20 seconds’, white regions in the merged image represent colocalization of signal). Scale bar – 10 μm. Reaction buffer contains 50 mM HEPES pH 7.3, 5 mM Mg(CH_3_COO)_2_, 300 mM KCH_3_COO, 50 mM KCl, 10% glucose, 0.1mg/mL casein (blocking agent).

To compare our reconstitution approach to previously published studies on FtsZ polymers on SLBs, we first examined structures formed by *E. coli* FtsZ with the YFP derivative venus and MTS fused to its C-terminus in tandem and replacing the CTC (*Ec* His_6_-FtsZ-venus-MTS) (Osawa *et al.*, 2008). This is modeled after *Ec* FtsZ-YFP-MTS which has been used in the past to observe *E. coli* FtsZ polymerization on membranes, within vesicles as well as on planar SLB (Osawa *et al.*, 2008; Osawa *et al.*, 2009; Osawa and Erickson, 2011; Arumugam *et al.*, 2012; Osawa and Erickson, 2013; Loose and Mitchison, 2014; Arumugam *et al.*, 2014; Ramirez *et al.*, 2016). When we flowed in 2 μM *Ec* His_6_-FtsZ-venus-MTS premixed with 2 mM GTP for 30 minutes, we observed dynamic assembly of fluorescent clusters on the membrane (Figure1B, Movie 1.1.). The clusters were typically amorphous or circular, measuring 413 nm ± 53 nm along short axis and 615 nm ± 94 nm along long axis (mean ± S.D., n = 28) at the start of imaging (Figure 1C, D). These measurements are likely overestimations of the actual dimensions of the structures due to the resolution-limit of light microscopy (~200 nm). Within 5 minutes of assembly, the amorphous clusters extended into dynamic filaments with variable lengths (980 ± 350 nm, mean ± S.D., n = 29) and relatively constant widths (381 nm ± 59 nm, mean ± S.D., n =29) (Figure 1B, 1C, 1D). On further incubation, these filaments arranged into regular patterns made of slightly curved filament bundles of similar widths as their precursors (425 nm ± 84 nm, mean ± S.D., n = 25) with periodic fluorescence intensity fluctuations along the length of the bundles (Figure 1D, 1E, Movie1.1). The bundles that constituted these structures appear similar in dimensions and dynamics to published observations of assembly on SLB-coated wells of *Ec* FtsZ-YFP-MTS (Arumugam *et al.*, 2014) or *Ec* FtsZ polymers with the membrane-anchoring protein ZipA (Loose and Mitchison, 2014).

When we flowed 1.8 μM *C. crescentus* FtsZ-venus-MTS preincubated with 2 mM GTP for 30 minutes into the flow cell, we observed the formation of amorphous or circular clusters (minimum width of 397 nm ± 62 nm, mean ± S.D., n = 26) similar to those observed at early time points with *Ec* His_6_-FtsZ-venus-MTS (Figure 1D, 1F, Movie 1.2). These clusters assembled into speckled patterns at steady state (Figure 1G, Movie 1.3) while their dimensions remained similar to their precursors (minimum width of 446 nm ±77 nm, mean ± S.D., n = 26) (Figure 1D). In contrast to the regular bundled patterns formed by *Ec* His_6_-FtsZ-venus-MTS that remained stable for minutes (Figure 1E), the *Cc* FtsZ-venus-MTS structures formed highly dynamic and irregular speckled patterns on the SLB surface (Figure 1H). We conclude that *C. crescentus* FtsZ assembles dynamically on SLBs into superstructures that are distinct from the filamentous superstructures formed by *E. coli* FtsZ.

### In addition to small dynamic clusters, *C. crescentus* ΔCTL forms very large multi-filament bundles on SLB

To address the contributions of the CTL to the assembly properties of *C. crescentus* FtsZ on membranes, we compared FtsZ with ΔCTL and other CTL variants. Since we are interested in the contributions of the unstructured C-terminal region of FtsZ, we decided against using the bulky fluorescent fusion (venus-MTS) at the C-terminus of our CTL variants. Instead, we used FtsZ or ΔCTL fluorescently labeled with Alexa488 dye conjugated at the only cysteine residue in *C. crescentus* FtsZ (Cys123) to visualize polymers. In addition to FtsZ or ΔCTL (a fraction of which was Alexa488-labeled), we also included equimolar unlabeled MTS-fusions of FtsZ or ΔCTL, wherein the CTC was replaced by the MTS, to recruit polymers to the membrane.

First, we confirmed that FtsZ-MTS could be used to specifically recruit FtsZ polymers to the membrane using a 1:1mixture of FtsZ and FtsZ-MTS (Figure 2A). On introducing 2 μM FtsZ (35% FtsZ-Alexa488) pre-incubated with 2 mM GTP for 5 minutes into a flow cell equilibrated with 2 μM FtsZ (35% FtsZ-Alexa488), we observed a minor increase in fluorescence intensity above background levels (Figure 2B, 2D, Supplementary figure 1A, Movie 2.1). This increase was accompanied by the appearance of dynamic fluorescent clusters (Figure 2B, 2D, Supplementary figure 1A, Movie 2.1) suggesting the formation of FtsZ polymers in the solution phase that can transiently contact the SLB surface. Strikingly, when we subsequently flowed in 2 μM FtsZ (35% FtsZ-Alexa488) and 2 μM FtsZ-MTS (unlabeled) pre-incubated with 2 mM GTP, we observed a rapid increase in the number and intensity of fluorescent clusters (Figure 2C, Movie 2.2, Supplementary figure 1A). Since FtsZ-MTS is not fluorescently labeled, we conclude that the increase in intensity is due to the co-polymerization of FtsZ and FtsZ-MTS at the membrane bringing Alexa488-labeled FtsZ into the TIRF illumination field. At steady state, the dynamic clusters organized into speckled cloud-like patterns with fluctuating local fluorescence intensities (Movie 2.3, Supplementary figure 1B). On photobleaching, the FtsZ/FtsZ-MTS structures took 23.7 s ± 1.9 s (mean ± S.D., n = 3) to recover half the maximum intensity (Supplementary figure 1C), confirming that these structures are undergoing rapid turnover.

**Figure 2.**
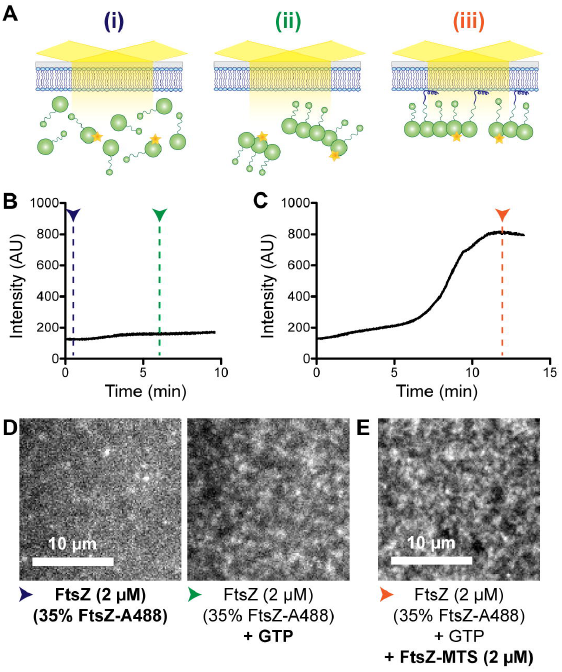
FtsZ-MTS co-polymerizes with FtsZ and recruits protofilaments to SLBs. **A.** Schematic corresponding to the experimental setup in B – E. **(i)**. Flow cell containing 20% DOPG and 80% DOPC SLB equilibrated with 2 μM FtsZ (35% FtsZ-Alexa488), **(ii)** At steady state after flowing in 2 μM FtsZ (35% FtsZ-Alexa488) with GTP, and **(iii)** At steady state after subsequently flowing in 2 μM FtsZ (35% FtsZ-Alexa488) and 2 μM FtsZ-MTS (unlabeled) with GTP. **B. & C.** Fluorescence intensity on the SLB averaged over the frame (~ 400 μm^2^) over time. **B.** 2 μM FtsZ (35% FtsZ-Alexa488) with GTP was flowed at 0.5 μL/minute into a flow cell equilibrated with 2 μM FtsZ (35% FtsZ-Alexa488). **C.** 2 μM FtsZ (35% FtsZ-Alexa488) and 2μM FtsZ-MTS with GTP was flowed at 0.5 μL/minute into a flow cell equilibrated with 2 μM FtsZ (35% FtsZ-Alexa488) with GTP. **D.** Contrast enhanced TIRFM images showing structures corresponding to experiment in B, immediately after beginning flow (blue arrowhead) and at steady state (green arrowhead). Contrast enhanced TIRFM image showing structures corresponding to experiment in C at steady state (orange arrowhead). Blue, green and orange arrowheads correspond to stages (i), (ii), and (iii) respectively as depicted in A. Scale bar – 10μm. Reaction buffer contains 50 mM HEPES pH 8.0, 0. 1mM EDTA, 2.5 mM MgCl_2_, 300 mM KCl, 1% glycerol, 0.1mg/mL casein (blocking agent), 0.5 mg/mL ascorbate.

Next, we turned our attention to the role of the CTL in regulating FtsZ assembly on SLB. Since we could not attain labeling efficiency greater than 6% for ΔCTL, we used 6% Alexa488 labeled FtsZ or ΔCTL in our experiments. On introduction of 2 mM GTP into a reaction containing 2 μM FtsZ (6% Alexa488 labeled) with 2 μM FtsZ-MTS, we observed structures similar to those observed for 2 μM FtsZ (35% Alexa488 labeled) with 2 μM FtsZ-MTS (Figure 3A, Movie 3.1). By measuring change in intensity following the introduction of FtsZ-MTS or GTP into flow cells equilibrated with FtsZ and GTP, or FtsZ and FtsZ-MTS, correspondingly, we confirmed that the assembly of these structures on the SLB was MTS- and GTP-dependent (Supplementary figure 2A, 2B).

**Figure 3.**
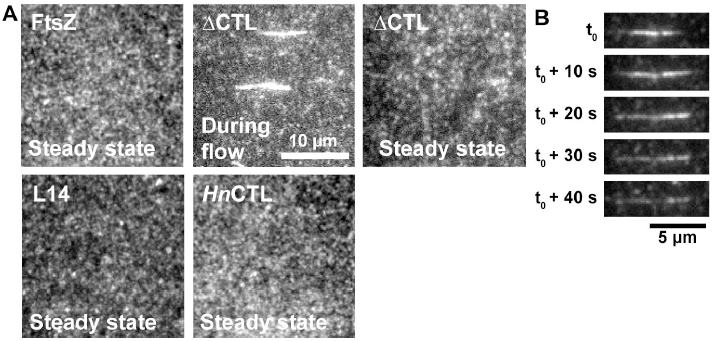
ΔCTL protofilaments form extended bright structures in addition to dynamic clusters. **A.** Contrast enhanced TIRFM images of structures observed on 20% DOPG 80% DOPC SLBs for FtsZ or CTL variants flowed in with 2 mM GTP at steady state or during flow. The FtsZ variants in each of the flow cells are 2 μM FtsZ or CTL variant (6% FtsZ-Alexa488 or corresponding Alexa488-labeled CTL variant) and 2 μM C-terminal MTS fusions replacing the CTC of FtsZ or corresponding CTL variant. **B.** Representative contrast enhanced TIRFM images showing the disassembly of an extended bright structure formed by ΔCTL/ΔCTL-MTS on the membrane over time. Scale bar – 10 μm. Reaction buffer contains 50 mM HEPES pH 8.0, 0.1.mM EDTA, 10 mM MgCl_2_, 300 mM KCl, 1% Glycerol, 01mg/mL Casein (blocking agent), 0.5 mg/mL ascorbate. Structures presented are representative and were confirmed using at least 3 independent replicates. Extended bright structures were observed in all replicates for ΔCTL/ΔCTL-MTS.

Intriguingly, with 2 μM ΔCTL (6% Alexa488 labeled) and 2 μM ΔCTL-MTS, we observed the rapid appearance of extended bright structures on the SLB following the introduction of GTP in addition to dynamic clusters similar to those observed for FtsZ/FtsZ-MTS (Figure 3A, 3B, Movie 3.2). Most of these structures oriented parallel to the direction of flow. After their rapid appearance, these structures disassembled gradually (Figure 3B, Supplementary figure 2C). At steady state, ΔCTL/ΔCTL-MTS protofilaments assembled as structures similar to FtsZ/FtsZ-MTS (Figure 3A, Supplementary figure 2D). However, these structures were comparatively sparse – whereas the local intensities of FtsZ/FtsZ-MTS patterns appear diffuse when averaged over time, ΔCTL/ΔCTL-MTS patterns contain more gaps between regions of high average intensities (Supplementary figure 2D). This result is in line with the sparse appearance of ΔCTL polymers on electron microscopy grids and the lower steady state light scatter observed for ΔCTL compared to WT FtsZ in solution (Sundararajan et al 2015, Sundararajan and Goley 2017).

The dimensions of the elongated structures of ΔCTL/ΔCTL-MTS on SLB are similar to the largest multi-filament bundles previously observed for ΔCTL polymers by electron microscopy (Sundararajan and Goley, 2017). Such bundles were never observed for WT FtsZ or CTL variants, namely L14 (FtsZ with a 14 amino acid CTL) and *Hn*CTL (FtsZ with CTL sequence from *Hyphomonas neptunium*) (Sundararajan and Goley, 2017). When we tested the assembly of L14/L14-MTS or *Hn*CTL/*Hn*CTL-MTS copolymers on SLBs, we did not observe any elongated structures. Similar to FtsZ/FtsZ-MTS assembly on membranes, L14/L14-MTS and *Hn*CTL/*Hn*CTL-MTS assembled into speckled cloud-like structures composed of dynamic fluorescent clusters at steady state (Figure 3A, Movie 3.3, Movie 3.4). From these observations, we conclude that the elongated structures observed specifically for ΔCTL/ΔCTL-MTS on SLB are large multi-filament bundles.

### *In situ* assembly/disassembly of FtsZ protofilaments on SLBs

While the appearance of large ΔCTL/ΔCTL-MTS bundles on SLBs is consistent with previous observations from electron microscopy that the CTL regulates lateral interaction between protofilaments (Sundararajan and Goley, 2017), we suspected that these structures are assembled in solution (upstream of the flow cell) during pre-incubation with GTP. Because we are interested in observing the behavior of structures that form on membranes *de novo*, we therefore altered the flow cell setup to rapidly control the availability of GTP within the flow cell allowing us to induce polymerization (or depolymerization) *in situ*. We flowed the protein mixture and GTP through two separate, parallel inputs into the flow cell with equal flow rates (Figure 4A). During flow, the protein and GTP channels meet within the flow cell and maintain a laminar boundary (Figure 4A). As long as flow is maintained, the laminar boundary acts as a diffusion barrier and constrains polymerization to the interface between the protein and GTP channels (Figure 4B-E, Movies 4.1 – 4.4). When flow is stopped, the two channels mix by diffusion, rapidly initiating polymerization on the protein side due to the much faster diffusion of GTP compared to FtsZ monomers or polymers. Restarting flow rapidly depletes GTP from the protein side, thereby favoring depolymerization and disassembly of FtsZ polymers. Thus, by controlling the flow, we can initiate assembly and disassembly of FtsZ polymers *in situ* within the flow cell (Figure 4A).

**Figure 4.**
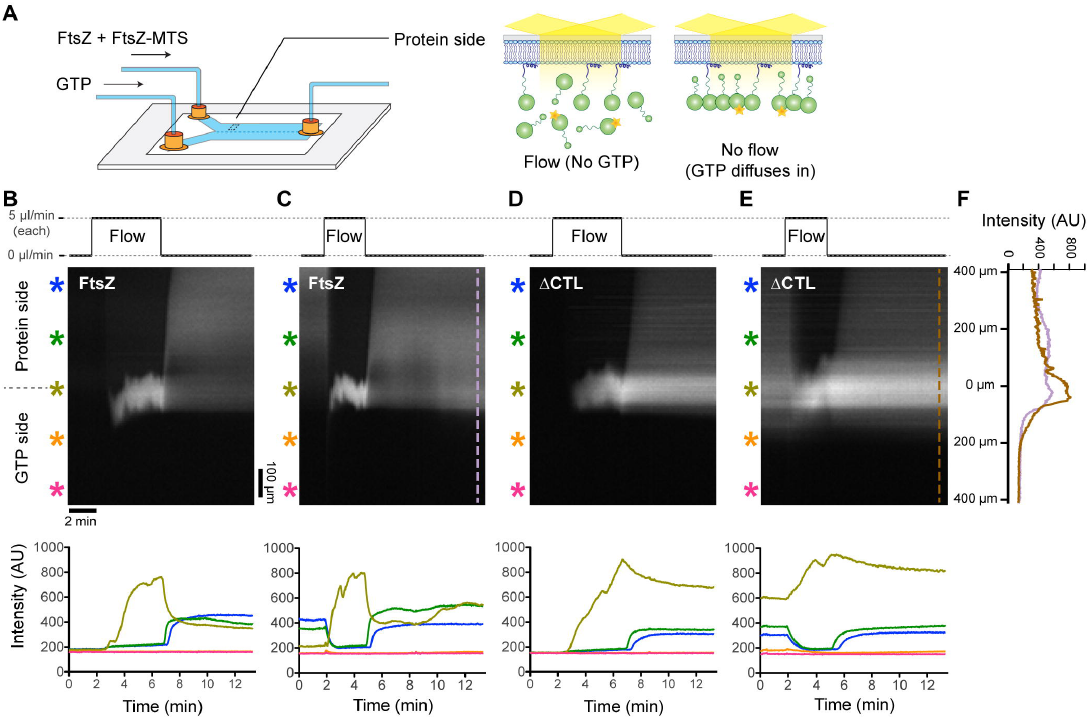
Flow-dependent control of assembly and disassembly of FtsZ and ΔCTL polymers. **A.** Schematic depicting the two-inlet flow cell used for rapid initiation of polymerization and depolymerization. During flow, the protein channel side is depleted of GTP and FtsZ is predominantly monomeric. Immediately after flow is stopped, GTP diffuses into the protein channel side initiating FtsZ (6% Alexa488 labeled) polymerization and recruitment to the membrane by copolymerizing with FtsZ-MTS, enabling visualization by TIRFM. **B-E**. Kymographs and corresponding fluorescence intensity vs time plots during periods of flow and no flow in the two-inlet flow cell. In the protein side, during flow, 2 μM FtsZ (6% Alexa488 labeled) and 2 μM FtsZ-MTS (unlabeled) (**B**, **C**) or 2 μM ΔCTL (6% Alexa488 labeled) and 2 μ ΔCTL-MTS (unlabeled) (**D, E**) is introduced at the flow rate of 5 μL/minute. Simultaneously, in the GTP side, 2 mM GTP is introduced at the same flow rate of 5 μL/minute. Time-lapse TIRF movies corresponding to the kymographs in B-E were obtained at 10x magnification. **B.** and **D.** represent kymographs and intensity plots corresponding to the first flow/stop cycle (flow up to 25 μL at 5 μL/minute for each channel into a fresh flow cell and then no flow to allow mixing), **C.** and **E.** correspond to subsequent flow-stop cycle (flow up to 15 μL at 5 μL/minute for each channel into the flow cell in B or D following steady state and then no flow to allow mixing).Scale bar = 100 μm in spatial axis (vertical) and 2 min in temporal axis (horizontal) of kymograph, asterisks of different colors correspond to intensity plots of the same color denoting regions within the flow cell at varying distances perpendicular to the laminar boundary (and the direction of flow). **F.** Line plots along axis perpendicular to the direction of flow at steady statefollowing re-initiation of assembly (after flow) at the indicated time points corresponding to kymographs in **C**. (FtsZ/FtsZ-MTS) and **E.** (ΔCTL/ΔCTL-MTS). Reaction buffer contains 50 mM HEPES pH 8.0, 0.1.mM EDTA, 10 mM MgCl_2_, 300 mM KCl, 1% glycerol, 0.1 mg/mL casein (blocking agent), 0.5 mg/mL ascorbate.

To confirm that we can achieve such flow-dependent control on FtsZ assembly, we imaged the microfluidic chamber at 10x magnification while simultaneously flowing 2 μM FtsZ (6% FtsZ-Alexa488) and 2 μM FtsZ-MTS mixture in the protein channel and 2 mM GTP in the GTP channel and subsequently stopping flow. Initially, on starting flow, we observed a rapid increase in fluorescence intensity only at the laminar boundary between the protein and GTP channels (Figure 4B, Movie 4.1). On the protein side, we observed a minor increase in fluorescence intensity likely due to unbound fluorescently labeled FtsZ monomers within the evanescent volume close to the SLB surface. On the GTP side, there was no significant increase in fluorescence intensity above background levels (Figure 4B, Movie 4.1). Immediately after the flow was stopped, we observed an increase in fluorescence intensity that spread gradually into the protein side, perpendicular to the original laminar boundary. On restarting flow, the average fluorescence intensity on the protein side decreased quickly until reaching levels comparable to background (Figure 4C, Movie 4.2). Subsequently, after the flow was stopped, the average fluorescence intensity increased once again, returning to values comparable to those observed before the flow was re-started (Figure 4C). The flow-dependent changes in fluorescence intensities are as expected for diffusion-limited introduction (flow, then stop), depletion (subsequent flow) and repletion (subsequent stop) of GTP in the protein side, and the corresponding induction of polymerization, depolymerization, and repolymerization of FtsZ/FtsZ-MTS copolymers on the SLB (Figure 4A).

The fluorescence intensity profiles over time were comparable between flow cells with FtsZ/FtsZ-MTS or ΔCTL/ΔCTL-MTS (Figures 4B – 4E), with two major differences. Firstly, at steady state (no flow), flow cells with FtsZ/FtsZ-MTS attained higher local fluorescence intensity values on the protein side and lower local fluorescence intensity values at the original laminar boundary compared to ΔCTL/ΔCTL-MTS (Figure 4F, Supplementary figure 3A). Secondly, fluorescence intensity in flow cells with ΔCTL/ΔCTL-MTS took significantly longer to drop back to background levels on restarting flow as discussed below (Supplementary figure 3B). These differences between FtsZ and ΔCTL intensity profiles observed at 10x magnification suggest that the CTL influences higher order assembly of FtsZ polymers on membrane.

### ΔCTL polymers assemble into relatively stable filament networks on SLB

Next, we observed the structures formed by FtsZ/FtsZ-MTS on the protein side of the original laminar boundary at 100x magnification. Immediately after stopping flow, we observed dynamic fluorescent clusters that assembled into speckled structures at steady state (Figure 5A, Figure 5B, Movie 5.1, 5.2) similar to our observations in the one-inlet flow cell setup. On restarting flow, these patterns gradually disassembled into sparse dynamic clusters that eventually disappear (Supplementary figure 3C, Movie 5.3). On stopping flow again, dynamic clusters reappear and form patterns distinct from those formed previously (before flow) (Supplementary figure 3C).

**Figure 5.**
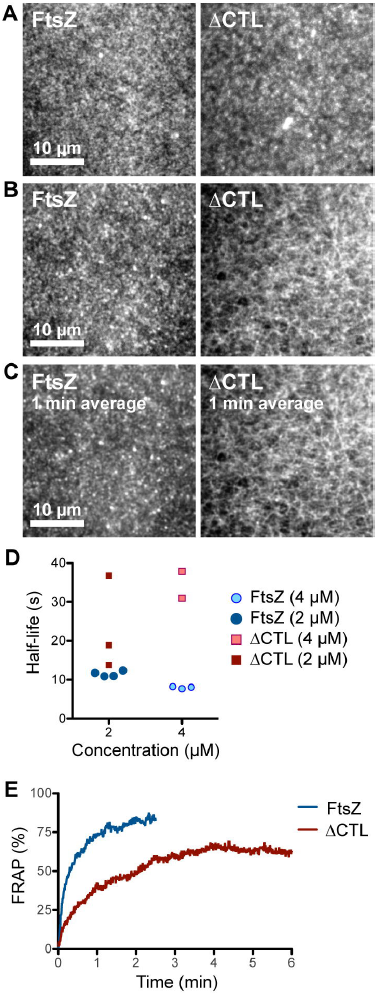
ΔCTL forms stable networks of straight filaments unlike WT FtsZ. **A, B.** Contrast enhanced micrographs of structures formed on SLBs at steady state after simultaneously flowing in 4 μM FtsZ (6% Alexa488 labeled) and 4 μM FtsZ-MTS (unlabeled) or 4 μM ΔCTL (6% Alexa488 labeled) and 4 μM ΔCTL-MTS (unlabeled) in the protein channel inlet and 4 mM GTP in the GTP channel inlet and stopping flow. **A.** Structures formed farther from the original laminar boundary on the protein side. **B.** Structures formed closest to the original laminar boundary on the protein side. **C.** Time averages corresponding to the structures shown in B. obtained from taking averages over frames corresponding to a 1-minute time interval. Scale bar– 10 μm. **D.** Time until decrease in fluorescence intensity to half-maximum value (half-life) for structures formed by FtsZ (6% Alexa488 labeled) and FtsZ-MTS (unlabeled) or ΔCTL (6% Alexa488 labeled) and ΔCTL-MTS (unlabeled), following depletion of GTP. Half-life values were estimated from non-linear fits assuming one-phase exponential decay. **E.** Fluorescence recovery after photobleaching corresponding to structures showed in A (on the protein side, away from the original laminar boundary). Plot shows average of 3 replicates. Reaction buffer contains 50 mM HEPES pH 8.0, 0. 1.mM EDTA, 10 mM MgCl_2_, 300 mM KCl, 1% glycerol, 0.1.mg/mL casein (blocking agent), 0.5 mg/mL ascorbate.

Strikingly, in addition to forming small dynamic clusters similar to those formed by FtsZ/FtsZ-MTS, ΔCTL/ΔCTL-MTS structures formed elongated filament bundles that interconnected into a stable network (Figure 5B, Movie 5.4, 5.5). While the fluorescence intensities within the network showed rapid fluctuations, the ΔCTL/ΔCTL-MTS network itself appeared stable for minutes (Figure 5C, Movie 5.4). These structures showed the highest fluorescence intensities in regions closest to the original laminar boundary on the protein channel side (Figure 4F). Unlike the 1-inlet setup, we did not observe thick, individual bundles of ΔCTL/ΔCTL-MTS copolymers on the SLB when polymerized *in situ*.

In addition to the structural differences between the polymers formed by FtsZ/FtsZ-MTS and ΔCTL/ΔCTL-MTS on SLB, we also observed significant differences in their dynamics. When we rapidly depleted GTP from the protein side by restarting flow, we observed that whereas the FtsZ structures disassembled at the rate of 3.6 ± 0.1 min^−1^ (mean ± S.D., n = 4, 2 μM total protein), ΔCTL/ΔCTL-MTS structures disassembled at a slower rate of 2. 1 ± 0.5 min^−1^ (mean ± S.D., n = 3, 2 μM total protein) (Figure 5D, Supplementary figure 3D, 3E). The decreased rate of disassembly of ΔCTL/ΔCTL-MTS on depleting GTP mirrored the decreased rate of fluorescence recovery after photobleaching observed for these structures compared to FtsZ/FtsZ-MTS (Figure 5E). Whereas FtsZ/FtsZ-MTS took 13 s ± 3 s (mean ± S.D., n = 3) to recover 50% of fluorescence following photobleaching, ΔCTL/ΔCTL-MTS took 42 s ± 8 s (mean ± S.D., n = 3 to recover fluorescence intensity with about 35% loss in fluorescence intensity following photobleaching. These results indicate that the structures formed by ΔCTL/ΔCTL-MTS are more stable and have slower turnover compared to FtsZ/FtsZ-MTS.

## Discussion

For polymerizing proteins such as FtsZ, their assembly properties are essential for their function. Observing the assembly of FtsZ polymerization in its physiological context is challenging in part due to the limitations of the spatio-temporal resolution of light microscopy and the complexity of multiple interacting components. *In vitro* reconstitution techniques have proven valuable for observing the assembly of dynamic cytoskeletal protein polymers from eukaryotes, and more recently, from bacteria. In the current study, we have described an *in vitro* reconstitution approach for observing FtsZ polymerization on planar SLBs, which, in addition to providing high spatial and temporal resolution, enables control of reaction conditions. As a validation of our approach, we demonstrate the reconstitution of His_6_-*Ec* FtsZ-venus-MTS polymers into dynamic patterns (Figure 1B) that are in agreement with the results of prior reconstitution efforts using *E. coli* FtsZ on SLBs (Arumugam *et al.*, 2012; Loose and Mitchison, 2014; Arumugam *et al.*, 2014; Ramirez *et al.*, 2016). Unlike for *E. coli* FtsZ or *Ec* His_6_-FtsZ-venus-MTS on SLB, we never observed stable patterns of dynamic filament bundles for *C. crescentus* FtsZ on SLBs. Instead, we observed dynamic clusters that organize into speckled patterns (Figure 1F). Using our approach to understand the effects of the CTL on regulating lateral interaction between *C. crescentus* FtsZ protofilaments, we observed that *C. crescentus* ΔCTL forms networks of straight filamentous structures (Figure 5A, B) similar in scale to the multi-filament bundles observed by electron microscopy (Sundararajan and Goley, 2017).Moreover, we observed significantly slower dynamics for the higher order assembly of ΔCTL protofilaments compared to FtsZ protofilaments (Figure 5D, 5E). Thus, our approach provides valuable insights into *C. crescentus* FtsZ polymerization in the context of the membrane and complements the previous biochemical characterization of the effects of the CTL.

Interestingly, the precursors to the superstructures formed by *Ec* His_6_-FtsZ-venus-MTS and *Cc* FtsZ-venus-MTS look comparable (Figure 1E). In both cases, dynamic clusters that are approximately 400 nm in diameter (or width) appear on the SLBs at the initial stage of polymer assembly. Similar dynamic clusters were observed with the CTL-variants of *Cc* FtsZ examined here, soon after the addition of GTP (Figure 3A). While the spatial resolution of the imaging system used here does not yield information on the organization within these nucleotide-dependent clusters, these clusters likely correspond to short individual protofilaments of FtsZ or bundles of a small number of short filaments, as observed by electron microscopy (Sundararajan and Goley, 2017). The assembly of *Ec* FtsZ and *Cc* FtsZ polymers into distinct dynamic superstructures despite the apparent similarity in their protofilament precursors is intriguing. Which, if any, of the superstructures formed by *Ec* FtsZ or *Cc* FtsZ on SLBs *in vitro* are relevant in the physiological context of Z-ring assembly? Individual clusters of FtsZ protofilaments in Z-rings *in vivo* are asymmetric and shorter than 200 nm in length as observed by electron cryotomography (Li *et al.*, 2007) or super-resolution light microscopy (Fu *et al.*, 2010; Holden *et al.*, 2014; Yang *et al.*, 2017), most similar to the precursors of *Cc* FtsZ or *Ec* FtsZ superstructures we observe here. In contrast, the emergent bundles of *Ec* FtsZ at steady state extend longer than 2 μm, dimensions not reported in cells for *E. coli* FtsZ. While the patterns formed by *E. coli* FtsZ protofilaments provide insights into the effects of constraining gently curved dynamic filaments to a flat and fluid surface (Ramirez *et al.*, 2016), their relevance to understanding FtsZ assembly *in vivo* might thus be limited.

An important variation in FtsZ across species is the length and sequence of the CTL (Vaughan *et al.*, 2004). Whereas *E. coli* FtsZ has a CTL of 48 amino acids, *C. crescentus* FtsZ has a much longer CTL of 172 amino acids. Curiously, when high concentrations of *B. subtilis* FtsZ CTL variants were polymerized in solution and observed by cryo-electron microscopy, the minimum spacing between adjacent protofilaments was found to correlate with the presence and length of the CTL (Huecas *et al.*, 2017). It is possible that the difference in the length and sequence of the CTL between *E. coli* FtsZ and *C. crescentus* FtsZ gives rise to the differences in their emergent structures on SLBs *in vitro* by altering intrinsic lateral and longitudinal interactions.

As demonstrated in this study and previous characterizations, the CTL plays an important role in preventing excess lateral interactions in *C. crescentus* (Sundararajan and Goley, 2017) and *E. coli* (Wang *et al.*, 1997). ΔCTL forms bundles in solution that can be observed on carbon-coated grids by electron microscopy (Sundararajan and Goley, 2017) or SLBs by TIRFM (Figure 3). In addition to elaborating on the structural differences previously observed by electron microscopy, we observe clear differences in dynamics between FtsZ and ΔCTL using the approach described here. Our *in vitro* measurements of FtsZ dynamics on the membrane suggest that intrinsic dynamics of *C. crescentus* FtsZ are comparable to those of FtsZ from *E. coli* and *B. subtilis*. The time to attain half-maximum FRAP of *C. crescentus* FtsZ we observe (~ 20s, Supplementary figure 1C) is similar to measurements of FRAP for *E. coli* FtsZ on supported lipid bilayers (~ 10 s (Arumugam *et al.*, 2014)), or *in vivo* (~30 s (Stricker *et al.*, 2002), ~ 10 s (Anderson *et al.*, 2004; Buss *et al.*, 2015)). Similar recovery rates have been observed for *B. subtilis* FtsZ *in vivo* (~10 s (Anderson *et al.*, 2004)).). ΔCTL polymers have a slower GTP hydrolysis rate (Sundararajan and Goley, 2017) and take proportionally longer to disassemble after GTP depletion (Figure 5D, Supplementary Figure 3D, 3E) or to recover after photobleaching (Figure 5E). These results confirm that the decrease in GTP hydrolysis rate observed for ΔCTL is directly linked to its turnover. Interestingly, mutants of FtsZ with similar (or reduced) GTP hydrolysis rates compared to ΔCTL do not cause envelope bulging and cell lysis *in vivo* (Sundararajan *et al.*, 2015) or affect polymer structure *in vitro* (Sundararajan and Goley, 2017). Therefore, we postulate that the combined effects of the CTL on organization and turnover of protofilaments contribute to ΔCTL’s lethal effects on cell wall metabolism *in vivo*.

Although the intrinsic assembly properties of FtsZ from *C. crescentus* on SLBs differ from those of FtsZ from *E. coli* as reported here, the structures formed by each in cells are remarkably similar. This implicates regulatory factors *in vivo* in modifying the assembly properties of FtsZ to generate a Z-ring with the appropriate dynamics and structure to effect division. The repertoire of Z-ring associated proteins that affect protofilament bundling and/or turnover is vast and varies across species (Gueiros-Filho and Losick, 2002; Mohammadi *et al.*, 2009; Goley *et al.*, 2010; Galli and Gerdes, 2011; Durand-Heredia *et al.*, 2012; Woldemeskel *et al.*, 2017; Lariviere *et al.*, 2018). For example, while FzlA, an essential protein that binds and assembles FtsZ filaments into helical bundles *in vitro*, is conserved in alpha-proteobacteria including *C. crescentus*, it is absent from other bacteria including *E. coli* (Goley *et al.*, 2010; Lariviere *et al.*, 2018). On the other hand, ZapC and ZapD, which induce bundling of *E. coli* FtsZ *in vitro* are not conserved in *C. crescentus* and other organisms. Moreover, while *E. coli* ZapA bundles *E. coli* FtsZ protofilaments (Low *et al.*, 2004; Small *et al.*, 2007; Mohammadi *et al.*, 2009), *C. crescentus* ZapA has no appreciable effects on *C. crescentus* FtsZ protofilaments *in vitro* (Woldemeskel *et al.*, 2017). Such differences in the availability and activity of FtsZ-bundling proteins could rectify the species-specific differences in the intrinsic higher order assembly of FtsZ we observe here to yield similar *in vivo* structures (Figure 1). Determining the effects of FtsZ-bundling proteins and other regulators of FtsZ assembly on the higher order assembly of protofilaments on SLBs will provide further insight into the regulation of Z-ring structure and dynamics *in vivo*.

Overall, our study provides the first characterization of polymer structure and dynamics on the membrane for FtsZ from a species other than *E. coli*. We have added spatio-temporal detail to the regulatory effects of the CTL on inter-filament interaction and turnover of *C. crescentus* FtsZ. While the current study uses an artificial membrane targeting sequence to constrain FtsZ polymerization to the membrane, expanding the study to include physiological membrane anchoring proteins such as FtsA and FzlC will be important future work. Furthermore, a large number of components of the division machinery dynamically interact with FtsZ, including those directly involved in peptidoglycan synthesis remodeling. The extension of the cell-free reconstitution system described here to investigate the interaction between FtsZ and the division machinery would greatly contribute to our understanding of the bacteria cell division process.

## Experimental Procedures

### Purification of proteins

*Ec* His_6_-FtsZ-venus-MTS was expressed for purification in *E. coli* Rosetta(DE3)pLysS cells using pET28C vector pEG658. All *C. crescentus* FtsZ variants (including *Cc*FtsZ-venus-MTS – pEG717, WT FtsZ – pMT219, FtsZ-MTS – pEG1295, ΔCTL – pEG681, ΔCTL-MTS – pEG1293, L14 – pEG723, L14-MTS – pEG1297, *Hn*CTL – pEG676, *Hn*CTL-MTS–pEG1296) used in this study were expressed for purification in *E. coli* Rosetta(DE3)pLysS cells using pET21a expression vectors (Supplementary Table 1). Nucleotide sequence information for previously unpublished plasmids are provided in Supplementary information. All FtsZ variants were purified using the previously published protocol for purifying *C. crescentus* FtsZ (Sundararajan *et al.*, 2015; Sundararajan and Goley, 2017). Cells were induced for expression of FtsZ variants for 3 hours at 37 °C at OD600 = 1.0 and pelleted following induction. The cell pellets were resuspended in lysis buffer (50 mM Tris-HCl pH 8.0, 50 mM KCl, 1 mM EDTA, 10% glycerol, DNase I, 1 mM β-mercaptoethanol, 2 mM PMSF with cOmplete mini, EDTA-free protease inhibitor tablet (Roche)), and lysed using lysozyme treatment (1 mg/mL) for 1 hour, followed by sonication to complete lysis. The lysate was then centrifuged at 6000xg for 30 minutes to remove cell debris and the filtered supernatant was applied to an anion exchange column (HiTrap Q HP 5 mL, GE Life Sciences). Fractions containing the FtsZ variant were eluted using a linear gradient of KCl and were pooled. The FtsZ variant was then precipitated from the eluate using ammonium sulfate (20-35 % saturation depending on the FtsZ variant) and confirmed using electrophoresis (SDS-PAGE) and Coomassie staining. The ammonium sulfate precipitate was resuspended in FtsZ storage buffer (50 mM HEPES-KOH pH 7.2, 0.1 mM EDTA, 50 mM KCl, 0-1. mM EDTA, 1mM β-mercaptoethanol, 10% glycerol) and purified further using size-exclusion chromatography (Superdex 200 10/300 GL, GE Life Sciences). The purified protein in FtsZ storage buffer was then snap frozen in liquid nitrogen and stored at −80 °C.

After purification, FtsZ, ΔCTL, L14 and HnCTL were subjected to Alexa488 dye labeling using Alexa Fluor 488 C5 Maleimide (ThermoFisher Scientific) reagent and the manufacturer’s protocol. Purified FtsZ or FtsZ variant was treated for 1 hour with a 10 times molar excess of DTT in FtsZ storage buffer to reduce the only cysteine residue in FtsZ, followed by incubation with at least 10 molar excess of Alexa Fluor 488 dye solution for 2 hours at room temperature or overnight at 4 C. Following incubation, a 20 times molar excess of β-mercaptoethanol was added to quench excess reagent in the reaction and the labeled protein was purified using size-exclusion chromatography (Superdex 200 10/300 GL, GE Life Sciences). The fluorescent fractions were pooled, concentrated and the stored at −80 °C. Prior to freezing, the labeling efficiency (as percentage labeled) was determined using absorption measurements. ΔCTL had the lowest labeling efficiency (6%) compared to other FtsZ variants. Hence, all experiments involving comparisons of ΔCTL to other FtsZ variants were performed with 6% labeled FtsZ variant in the final reaction.

### Preparation of flow cells

One- and two-inlet flow cells were prepared as described previously with a few modifications (Vecchiarelli *et al.*, 2016). Quartz glass slides with drilled one or two inlet holes and one outlet hole each (Esco products) were cleaned by washing overnight in NOCHROMIX glass cleaner (Sigma), rinsed with ultrapure water, air dried, and treated with low-power plasma cleaning in the presence of argon and oxygen. A rectangular piece of 25-μm thick acrylic transfer tape (3M) of ~ 5 cm x ~ 3.5 cm was cut to demarcate the required chamber dimensions (for one-inlet flow cell, rectangular region of 4 mm wide x 3 cm long was cut out, for two-inlet flow cell, y-shaped region with a uniform width of 4 mm was cut out). The tape was placed between the glass slide and cover slip. Nanoports (Upchurch) adapters were attached to the slides above the holes with optical adhesive. The flow cell was then baked at 65°C for 1 hour.

We often observed that FtsZ protofilaments were preferentially recruited or excluded along parallel straight lines on the SLBs. We hypothesized that this was due to scratches along the glass surface, giving rise to extended regions of curved membrane. While we observed these ordered linear patterns for *Ec* His_6_-FtsZ-venus-MTS, *Cc* FtsZ-venus-MTS as well as with Alexa488-labeled FtsZ, they were most obvious in experiments using partially labeled FtsZ (for example, FtsZ (35% FtsZ-Alexa488)/FtsZ-MTS). To avoid loss in signal-to-noise in imaging regions adjacent to scratches and to prevent possible artifacts, we treated the glass slides with hydrofluoric acid (HF) to remove scratches on the surface in all our experiments that involved Alexa-labeled FtsZ variant. Glass slides were incubated in 20% HF solution for 2 minutes, and then washed by immersing in 100 mM CaCl_2_ solution bath, and rinsed well with water prior to wash with NOCHROMIX. HF treatment of glass slides eliminated the appearance of parallel straight lines.

### Preparation of SUVs

Minimum synthetic lipid mixtures were made using 33:67 or 20:80 combinations of 1,2-dioleoyl-sn-glycero-3-[phospho-rac-(1-glycerol)] (DOPG; Cat. No. 840475, Avanti) and 1,2-dioleoyl-sn-glycero-3-phosphocholine (DOPC; Cat. No. 850375, Avanti). The purchased synthetic lipids resuspended in chloroform at 25 mg/mL were mixed to appropriate ratios in glass tubes pre-rinsed with chloroform. After thorough mixing, the lipid mixture was dried by evaporating chloroform using dry N_2_ gas with constant rotation to make a thin layer of dry lipids and was dried further in a SpeedVac Concentrator (Savant) for 1 hour at 42 °C initially and 1 hour at room temperature subsequently. The dried lipid mixture was resuspended by vortexing in degassed TK150 buffer (25 mM Tris-HCl, pH 7.4, 150 mM KCl) to a lipid concentration of 5 mg/mL and was incubated overnight in the dark at room temperature in an N_2_ atmosphere (N2 box). The next day, the aqueous resuspension of lipids was mixed thoroughly by vortexing and was transferred to polystyrene tubes. The resuspension was sonicated at 23 °C immersed in a water bath sonicator (Qsonica model #Q700A) at 70 W for 5 minutes (30 s/pulse with 10 s rest) until the turbid resuspension (made of multilamellar vesicles of non-uniform dimensions) turned translucent and blue-shifted (corresponding to ~ 100 nm small unilamellar vesicle or SUVs). Under N_2_ atmosphere, the sonicate was then filtered using 0.2 micron filter to purify SUVs, aliquoted and stored in Teflon-coated and parafilm-sealed glass vials at 4 °C. SUV stocks were used within 5 weeks from the date of preparation.

### Preparation of SLBs

Supported lipid bilayers were made by triggering attachment of SUVs to plasma cleaned glass slide surface within the flow cell by incubation with 5 mM MgCl_2_ in TK150 buffer for 1 hour at 37 °C. The flow cell was first equilibrated by flowing in TK150 buffer pH 7.4 containing 5 mM MgCl_2_ (TK150M5). The SUVs from the stock solution were diluted to 0.5 mg/mL in TK150M5 buffer and the solution was incubated at 37° C for 5 minutes. 300 μL of the SUV solution in TK150M5 was then flowed in at 10 μL/min into the flow cell maintained at 37° C. The flow cell was then incubated for 1 hour at 37 ° C to allow fusion of SUVs to form supported lipid bilayers. The excess SUVs were removed by flowing in 500 μL of TK150M5 buffer. The flow cells with SLBs were then equilibrated for subsequent experiments by flowing in appropriate reaction buffers. The flow cells were maintained at 37 °C until mounting on the microscope stage and were maintained above 24 °C during experiments to maintain membrane fluidity by avoiding phase transition of SLBs at lower temperatures.

### FtsZ polymerization reactions

Imaging experiments involving *Ec* His_6_-FtsZ-venus-MTS or *Cc* FtsZ-venus-MTS were performed in HMKKG FtsZ polymerization buffer (50 mM HEPES-KOH pH 7.2, 5 mM MgCl_2_, 150 mM KCl, 50 mM K(CH_3_CO_2_) 10% glycerol) containing 1% casein (w/v) and 0.5 mg/mL ascorbate using 2 μM FtsZ-venus-MTS incubated with 2 mM GTP for 30 minutes prior to flowing into flow cells with SLBs made from 33% DOPG, 67% DOPC SUVs.

Imaging experiments involving FtsZ/FtsZ-MTS, ΔCTL/ΔCTL-MTS, L14/L14-MTS or HnCTL/HnCTL-MTS were performed in HEK300 FtsZ polymerization buffer containing 50 mM HEPES-KOH pH 8.0, 0. 1 mM EDTA, 10 mM MgCl_2_ (unless otherwise mentioned), 300 mM KCl with 1% casein (w/v) and 0.5 mg/mL ascorbate incubated with 2 mM GTP for 5 minutes as required prior to flowing into the flow cells with SLBs made from 20% DOPG, 80% DOPC SUVs. These reaction conditions and membrane composition were determined to be optimum for reducing non-specific interaction of FtsZ polymers with the membrane in the absence of the MTS to improve signal-to-noise ratio. The protein mixtures were filtered using centrifugal filters prior to addition of nucleotide to remove non-specific protein aggregates on SLBs.

### TIRF microscopy, imaging and analysis

Illumination and imaging were performed using instrumentation described previously (Vecchiarelli *et al.*, 2016). All TIRFM experiments were performed on flow cell mounted on an Eclipse TE200E microscope (Nikon) with a prism placed on top of the glass slide (with oil, NA =149, between prism and glass slide) and imaged through the coverslip (bottom) through Plan Apo 10X (NA = 0.45, air) or Plan Apo 100X (NA =1 4, oil immersed) objectives (Nikon). An Andor DU-879E camera was used for image acquisition with the following settings: digitizer – 3 MHz (14 bit-gray scale), preamplifier gain – 5.2, vertical shift speed, 2 MHz, vertical clock range – normal, electron-multiplying gain – 40, EM CCD temperature – –98 °C, baseline clamp – ON, exposure time100 ms.

The excitation at 488 nm for FtsZ-venus-MTS and Alexa fluor 488 labeled FtsZ was provided using a 488 nm diode-pumped solid-state laser (Sapphire, Coherent) at 8 μW. TIRF illumination had a Gaussian shape in the field of view that could be broadened using a diffuser at the incident beam. Images were acquired in regions of uniform illumination profile to improve signal to noise.

Images were acquired at 0.5, 2 or 5 seconds per frame as mentioned in movie legends using Metamorph 7 (Molecular Devices) to make time-lapse movies in ImageJ (National Institute of Health). Movies were made from 150 px × 150 px or 200 px × 200 px regions of interest (ROIs) that were cropped from 512 px × 512 px fields of view and brightness/contrast adjusted, by enhancing contrast by saturating the highest 2% of intensities for each frame. The same brightness adjustment was applied to each frame. Movies were then converted to Audio Video Interleave format (.avi). Representative still images were made from 5 s time averages at specified time points (i.e. 10 frame time average for 0.5 seconds per frame acquisition, 5 frame time average for 2 seconds per frame acquisition, and 2 frame time average for 5 seconds per frame acquisition), to improve signal to noise.

Dimensions of fluorescent clusters of *E. coli* His_6_-FtsZ-venus-MTS and *C. crescentus* FtsZ-venus-MTS and of bundles of *E. coli* His_6_-FtsZ-venus-MTS on SLBs in Figures 1D and 1E were estimated using line-scans across the short axis (width) or long axis (length) of these structures. The short and long axes were obvious mainly for *E. coli* His_6_-FtsZ-venus-MTS after 5 minutes on the SLB (Figure 1D). For circular or amorphous clusters, the shortest distance across the cluster was estimated. Fluorescent profiles were measured along lines drawn through the structures. The distances between points of half-maximum intensity (full width at half-maximum) were determined from polynomial fits to the fluorescence profiles that were generated using Graphpad Prism Software (Graphpad Software Inc., La Jolla, CA).

Intensity plot profiles were measured as averages of fluorescence intensities in regions of interest (entire frame –150 px × 150 px, 200 px × 200 px, FRAP – 40 px × 40 px ROI within photobleached region, ΔCTL bundles – minimum rectangular ROIs, approximately 8 px × 15 px, around bright filamentous structures) per frame.

Movies showing flow-stop specific intensity changes in two-inlet flow setup in figure 4 were acquired using a 10X objective. Corresponding kymographs were obtained at line (spline width 4 px) perpendicular to the direction of flow. 30 px × 30 px ROIs were used for measuring corresponding fluorescence intensity profiles over time in these experiments.

### GTP depletion experiments

Rapid GTP depletion to induce disassembly of FtsZ or ΔCTL polymers on SLBs were performed using the two-inlet setup as described in Figure 4. The rate of disassembly and half-lives of the polymers on SLBs after GTP depletion (Figure 5D) were estimated using exponential decay fits to fluorescence intensity profiles over time by TIRFM at 100x magnification averaged over ROIs of 40 px × 40 px ROI.

### Photobleaching experiments

Fluorescence recovery after photobleaching (FRAP) experiments were performed using high power laser applied for 3 seconds on the SLBs through the objective lens using ~ 6 times the intensity used for the incident light for TIRF, while momentarily pausing image acquisition. We observe a minimum 40% loss in fluorescence immediately following photobleaching in our FRAP experiments. Time to half-maximum FRAP were estimated using one-phase association curves fit to fluorescence intensity profiles of individual replicates.

## Acknowledgements

We would like to thank the members of the Goley lab – Elizabeth Meier, PJ Lariviere, Selam Woldemeskel, Anant Bhargava, Allison Daitch, and Chris Mahone – and the Mizuuchi lab – James Taylor and Michiyo Mizuuchi – for helpful discussions that informed design, optimization and analyses described in this work. We would also like to thank Jie Xiao, Xinxing Yang, and Keir Neuman for their suggestions for quantifying FtsZ dynamics on SLBs. Funding for this work was provided by the NIH through R01GM108640 (to E.D.G.) and the intramural research fund for National Institute of Diabetes and Digestive and Kidney Diseases (to K.M).

## Author Contributions

KS, AV, KM, and EDG designed the experiments. KS and AV performed the experiments. KS analyzed the data. KS, AV, KM, and EDG wrote the paper and approved the final version of the manuscript.

## Conflict of Interest

The authors declare they have no conflict of interests with the contents of this manuscript.

## Movie legends

**Movie 1.1:** *Ec* His_6_-FtsZ-venus-MTS protofilaments assemble into dynamic bundles on SLBs. Contrast enhanced time-lapse movie of 2 μM *Ec* His_6_-FtsZ-venus-MTS with 2 mM GTP introduced into the flow cell and allowed to assemble on SLB membrane made of 33% DOPG and 67% DOPC acquired at 5 frames per second. Scale bar – 10 μm. Speed – 20x.

**Movie 1.2:** *Cc* FtsZ-venus-MTS protofilaments assemble into resolution-limited spots on SLB. Contrast enhanced time-lapse movie of 1.8 μM *Cc* FtsZ-venus-MTS with 2 mM GTP introduced into the flow cell and allowed to assemble on SLB membrane made of 33% DOPG and 67% DOPC acquired at 2 frames per second. Scale bar – 10 μm. Speed – 20x.

**Movie 1.3:** *Cc* FtsZ-venus-MTS assembles into resolution-limited dynamic clusters at steady state. Contrast enhanced time-lapse movie of assembly of 1.8 μM *Cc* FtsZ-venus-MTS with 2 mM GTP on SLB membrane made of 33% DOPG and 67% DOPC acquired at 2 frames per second. Scale bar –10 μm. Speed – 20x.

**Movie 2.1:** FtsZ protofilaments form transient dynamic clusters on the SLB. Contrast enhanced time-lapse movie of 2 μM FtsZ (35% Alexa488 labeled) with 2 mM GTP flowed (at 0.5μL/minute) into flow cell equilibrated with 2 μM FtsZ (35% Alexa488 labeled) without GTP, onto SLB membrane made of 20% DOPG and 80% DOPC acquired at 0.5 frames per second. Scale bar –10 μm. Speed – 20x. (Representative of 3 replicates)

**Movie 2.2 & 2.3:** FtsZ/FtsZ-MTS protofilaments assemble into dynamic clusters. Contrast enhanced time-lapse movie of structures formed by 2 μM FtsZ (35% Alexa488 labeled) and 2μM FtsZ-MTS (unlabeled) with 2 mM GTP flowed (at 0.5 μL/minute) into flow cell equilibrated with 2 μM FtsZ (35% Alexa488 labeled) with 2 mM GTP without FtsZ-MTS, onto SLB membrane made of 20% DOPG and 80% DOPC acquired at 0.5 frames per second. Movie **2.2** corresponds to 0 – 5 minutes, Movie **2.3** corresponds to 5 minutes –13.5 minutes of experiment described in Figure 2C. Scale bar –10 μm. Speed – 20x. (Representative of 3 replicates)

**Movies 3.1 – 3.4:** CTL regulates FtsZ polymer structure on SLBs. Contrast enhanced time-lapse movies of structures formed by FtsZ variants with 2 mM GTP flowed into flow cell equilibrated with FtsZ variants alone without GTP, onto SLB membrane made of 20% DOPG and 80% DOPC acquired at 0.5 frames per second. Scale bar –10 μm. Speed – 20x. FtsZ variants in each flow cell correspond to 2 μM FtsZ or CTL variant (6% Alexa488 labeled) and 2μM FtsZ-MTS or MTS fusion to corresponding CTL variant (unlabeled). 1– FtsZ, **2** -ΔCTL, **3** – L14, and **4** – *Hn*CTL. (Representative of 3 replicates)

**Movies 4.1 – 4.4:** Flow-dependent rapid initiation of assembly and disassembly of FtsZ or ΔCTL polymers on SLB. Contrast enhanced time-lapse movies of structures formed by 2 μM FtsZ (1& **2**) or ΔCTL (**3 & 4**) (6% Alexa488 labeled) and 2 μM unlabeled FtsZ-MTS or ΔCTL-MTS, correspondingly, with 2 mM GTP at steady state on SLB membrane made of 20% DOPG and 80% DOPC acquired at 0.5 frames per second.1**,3** 25 μL of each input was flowed in simultaneously at 5 μL/minute into a flow cell equilibrated with buffer alone. **2,4** 15 μL of each input was flowed in simultaneously at 5 μL/minute into a flow cell following steady state in 1 and **3** correspondingly. Movies were acquired at 10x magnification at 2 seconds per frame. Scale bar –100 μm. Speed – 80x.

**Movie 5.1:** Initial assembly of FtsZ on SLBs in two-inlet flow cell setup. Contrast enhanced time-lapse movies of structures formed by 4 μM FtsZ (6% Alexa488 labeled) and 4 μM unlabeled FtsZ-MTS with 4 mM GTP at steady state on SLB membrane made of 20% DOPG and 80% DOPC acquired at 0.5 frames per second on the protein side after stopping flow. Scale bar –10μm. Speed – 20x. (Representative of at least 3 replicates)

**Movie 5.2:** FtsZ assembly on SLBs in two-inlet flow cell setup at steady state. Contrast enhanced time-lapse movies of structures formed by 4 μM FtsZ (6% Alexa488 labeled) and 4μM unlabeled FtsZ-MTS with 4 mM GTP at steady state on SLB membrane made of 20% DOPG and 80% DOPC acquired at 0.5 frames per second on the protein side close to the original laminar boundary after stopping flow. Scale bar –10 μm. Speed – 20x. (Representative of at least 3 replicates)

**Movie 5.3:** FtsZ polymers disassemble and reassemble on depletion and repletion of GTP. Contrast enhanced time-lapse movies of structures formed by 2 μM FtsZ (6% Alexa488 labeled) and 2 μM unlabeled FtsZ-MTS with 2 mM GTP at steady state on SLB membrane made of 20% DOPG and 80% DOPC acquired at 0.5 frames per second on the protein side showing dynamics during steady state (0:00 – 0:30), during flow (0:30 – 3:30) and after flow. 15 μL of each input was flowed in simultaneously at 5 μL/minute into flow cell following steady state. Scale bar –10 μm. Speed – 20x. (Representative of at least 3 replicates)

**Movie 5.4:** Initial assembly of ΔCTL on SLBs in two-inlet flow cell setup. Contrast enhanced time-lapse movies of structures formed by 4 μM ΔCTL (6% Alexa488 labeled) and 4 μM unlabeled ΔCTL-MTS with 4 mM GTP at steady state on SLB membrane made of 20% DOPG and 80% DOPC acquired at 0.5 frames per second on the protein side after stopping flow. Scale bar –10 μm. Speed – 20x. (Representative of at least 3 replicates)

**Movie 5.5:** ΔCTL forms stable networks of straight filament bundles unlike WT. Contrast enhanced time-lapse movies of structures formed by 4 μM ΔCTL (6% Alexa488 labeled) and 4μM unlabeled ΔCTL-MTS with 4 mM GTP at steady state on SLB membrane made of 20% DOPG and 80% DOPC acquired at 0.5 frames per second on the protein side close to the original laminar boundary after stopping flow. Scale bar –10 μm. Speed – 20x. (Representative of at least 3 replicates)

